# Single-aperture SLAM microscopy with amplitude-tailored vector beams

**DOI:** 10.1101/2025.10.07.679791

**Authors:** Jeffrey Demas, Iván Coto Hernández, J. Matthew Dubach, Siddharth Ramachandran

## Abstract

Switching laser mode (SLAM) microscopy is a promising method for achieving super resolution while maintaining compatibility with two photon imaging at depth and *in vivo*. SLAM microscopes typically employ multiple paths for generating the requisite spot-like and donut-like beams; however, having two paths necessitates sub-wavelength-scale alignment which is prone to differential drift, causing degradation of the image quality. Here we demonstrate a single aperture, inline SLAM microscope which makes use of one phase element and polarization switching to generate colinear radially polarized and azimuthally polarized vector beams, which focus to a spot and a donut, respectively. By tailoring the spatial profile of the electric field at the back aperture of the microscope objective, we ensure that the resolution of the spot-like beam is comparable to conventional Gaussian beam imaging. Through subtraction of the two images, we demonstrate a 1.5× narrower focal spot and a resolution of ∼0.28*λ* corresponding to ∼290 nm. Accordingly, this method is of great utility for imaging with sub-diffraction-limited resolution at depth in living tissue.

## 1. Introduction

In the past few decades, resolution in optical microscopy has far surpassed the limit posed by the diffraction of light, with a host of super-resolution imaging techniques reaching ∼nm-scale resolution (1–5). However, few of these technologies penetrate beyond the first ∼100 μm of tissue as they really on spatially coherent imaging of fluorescence photons, which is impractical due to scattering. Two photon microscopy (2PM) is the gold standard for imaging at depth as both the excitation and the detection of fluorescence photons are resilient to scattering (6); allowing for penetration as deep as 4-5 scattering lengths in tissue. The primary drawback, however, is that 2PM necessitates near-infrared excitation wavelengths, which reduces resolution. Even a modest increase in 2PM resolution to ∼100–200 nm – comparable to conventional, depth-limited techniques such as confocal or widefield imaging – would provide valuable additional information for myriad imaging applications: for example, tracking the motility, morphology, and activity of dendritic spines in the developing mammalian brain (7,8), or non-invasive imaging of peripheral nerves and corneal tissue (9,10).

Switching laser mode (SLAM) microscopy (11), also known as fluorescence emission depletion (FED) microscopy (12), provides both super-resolution and compatibility with two-photon excitation. In SLAM, two images are captured – one with a conventional Gaussian-spot-like point-spread function (PSF), and the other with an annular or “donut”-shaped beam; by subtracting the images, resolution can be improved relative to the diffraction limit. Developments in SLAM microscopy have employed many different variations in donut generation (13,14) and switching mechanisms (15), as well as improvements in post-processing to reduce artifacts (16,17) and improve resolution further (18). Currently, SLAM with two-photon excitation can enhance resolution by 1.4–1.6× relative to diffraction-limited imaging (11, 16).

Conventional SLAM microscopes employ two distinct paths for each imaging beam (Fig. 1a), with one path including a mode conversion element for creating the donut beam (11,16,19). Using two paths is an issue for long-duration imaging experiments as differential drift between the paths can offset the registration of the beams (20,21), causing artifacts and reduction in resolution. Here, we develop a SLAM microscope with a single aperture, inline path for each beam – eliminating the possibility of any differential drift (Fig. 1b). The microscope employs a vortex waveplate (VWP) which generates azimuthally and radially polarized cylindrical vector beams for incident vertical and horizontal polarization states, respectively.

**Fig. 1:**
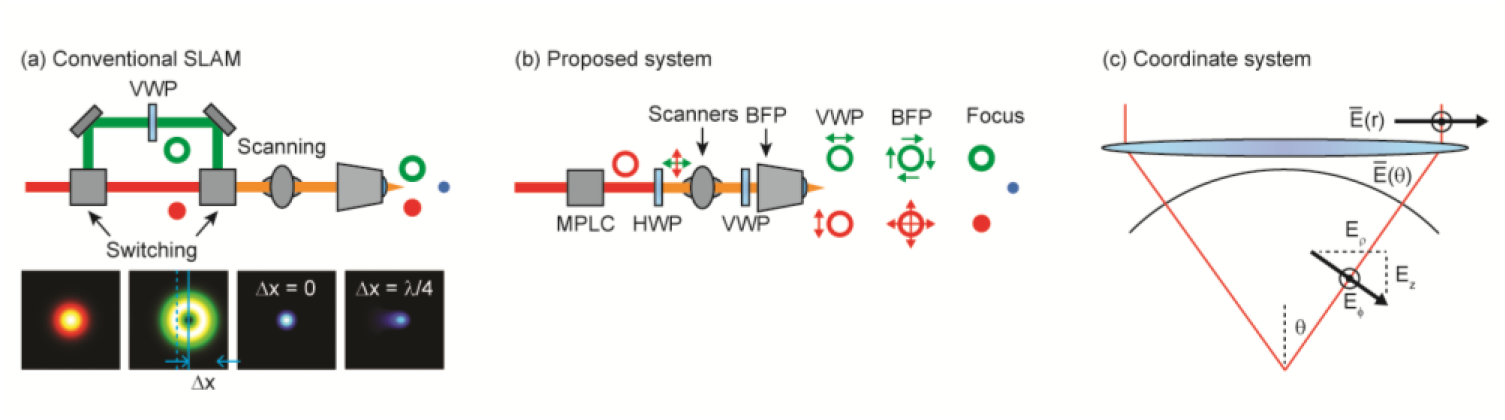
(a) Conventional SLAM microscopes employ switching (polarization- or mechanical-based) between two paths, allowing for differential drift (*Δx*) and degradation of the resulting PSF. (b) Schematic of a single aperture SLAM microscope: MPLC shapes the far field beam into a ring of either vertical or horizontal polarization; a VWP shapes the beam into an AP or RP beam at the BFP according to the incident polarization state; at focus, the AP beam remains a donut, while the RP beam becomes a spot facilitating resolution enhancement by subtraction; note that all beams have the same wavelength and different colors delineate different beam paths and/or polarizations; HWP = half-wave plate. (c) Transformation between the far field cylindrical coordinate system and the spherical near field by an aplanatic lens.

This scheme exploits the well-known fact that radially polarized (RP) beams generate a longitudinal polarization component under high NA focusing which collapses to a small focal spot (22–26). We employ a multi-plane light conversion (MPLC) system using a spatial light modulator (SLM) to generate tailored ring-shaped beams at the back focal plane (BFP) of the objective; by optimizing the aspect ratio of the ring, we increase the longitudinal component at focus to ensure a tight focal spot for the RP beam, providing resolution comparable to that of conventional Gaussian beam imaging. Conversely, the azimuthally polarized (AP) beam remains a donut under high NA focusing; thus, switching the linear polarization state incident on the VWP generates each of the requisite beam shapes at focus and enables two-photon SLAM imaging with a single path and aperture. Using multiple resolution criteria, we determine that the subtracted images demonstrate resolution as fine as *d* ∼ 0.28*λ* (290 nm) – a factor of ∼1.4× improvement in resolution relative to imaging with the RP beam alone – making this system attractive for imaging sub-cellular features in biological tissues.

## 1. Cylindrical vector beams at focus

In the scalar approximation, the polarization composition of an electric field is unchanged by focusing. However, for the non-paraxial case (*i*.*e*. high NA focusing), vector diffraction theory is required. Consider an electric field 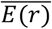 composed of RP and AP components, 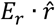 and 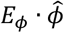, incident on an aplanatic lens (Fig. 1c) where *r* and *ϕ* are the far field radial and azimuthal coordinates, respectively. Mapping the vector field onto the reference sphere of the lens converts a portion of the RP component to a longitudinally polarized, or z-polarized (ZP) field (22–26). At focus, this ZP component forms a tight focal spot, whereas the residual RP light must remain a donut due to the polarization singularity at the center of the field. AP light is unchanged by the mapping process and similarly generates a donut at focus. This simple model also illustrates that the parts of the field closest to the periphery of the lens (*i*.*e*. large polar angle, *θ*) will have the most radial to longitudinal conversion. For this reason, illuminating the BFP of the objective with an annular RP light beam can be used to increase the ZP content at focus relative to the residual RP field, leading to spot-like foci (23).

To quantitively evaluate the effects of the amplitude structure of the far field beam on the spatial characteristics of the beam at focus, we calculate the vector diffraction integrals following the treatment of Richards and Wolf (27). We define the electric field incident on the BFP of the objective (Fig. 2a) as

**Fig. 2:**
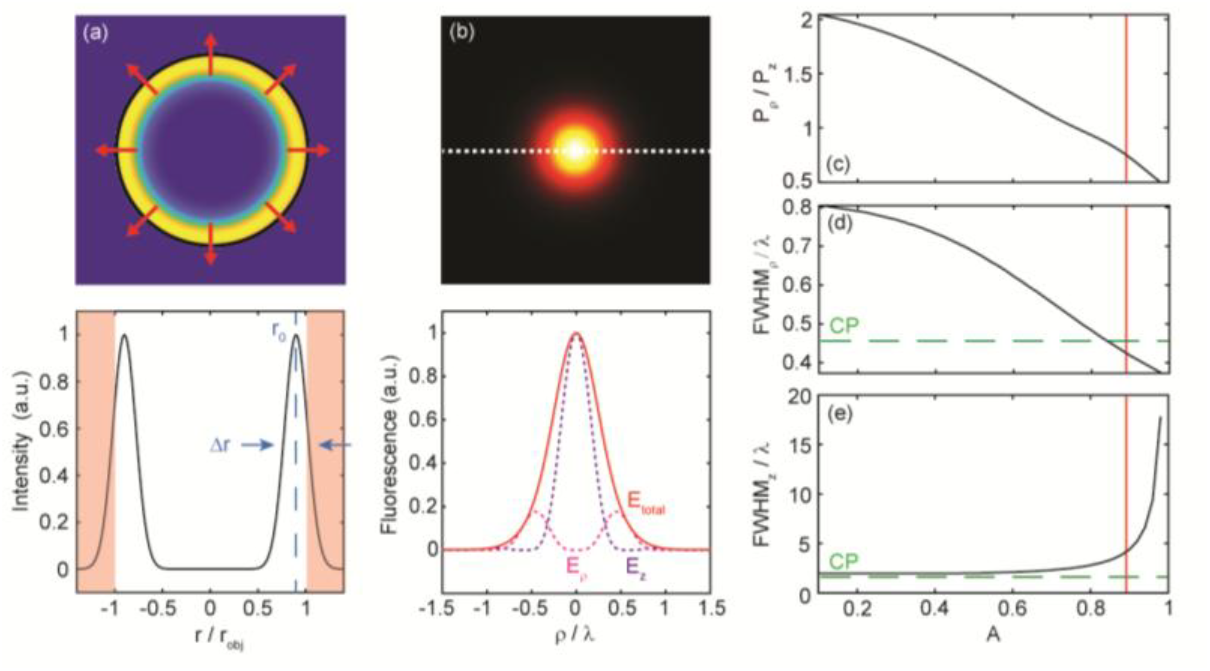
(a) Annular RP beam at the back focal plane of the objective (line cut below); red regions denote light cut off by the microscope back aperture; values *r*_*0*_ and *Δr* are parameterized by the aspect ratio (*A*) of the beam. (b) Focused RP beam generates a spot-like point spread function, with fluorescence contributions from both the converted longitudinal (*E*_*z*_, dashed purple line) component of the field and residual radially polarized (*E*_*ρ*_, dashed pink line) component (total field, *E*_*total*_, shown in solid red line). Ratio of the power in the radial (*P*_*r*_) and longitudinal components (*P*_*z*_) at focus (c), and lateral (d) and axial (e) full-width-at-half maximum (FWHM) versus aspect ratio; aspect ratio for this work (*A* = 0.9) marked with red line; green dashed lines denote values for a circularly polarized (CP) Gaussian beam at focus.

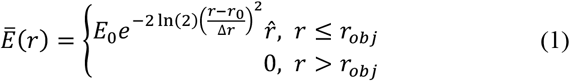

where *E*_*0*_ is the amplitude of the field. The electric field has the form of a Gaussian beam shifted to a center radius *r*_*0*_, with a full-width-at-half-maximum (FWHM) *Δr*. It is convenient to define these beam parameters in terms of a single value *A*, representing the aspect ratio of the ring, such that *r*_0_ = *r*_*obj*_ (1 + *A*)/2 and Δ*r* = *r*_*obj*_ (1 − *A*). Outside the radius of the back aperture, *r*_*obj*_, the electric field is zero. The far field beam is mapped to the reference sphere by the following relation

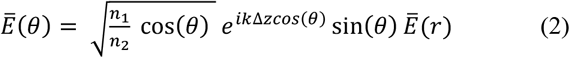

where *θ* is the polar angle, *n*_*1*_ and *n*_*2*_ are the refractive indices of the object and image environments, *Δz* is the distance from the focal plane, and *k* is the wave number given by *k* = 2*π*/*λ*, where *λ* is the wavelength of light. Due to the cylindrical symmetry of the problem, trigonometric identities can be employed to express the diffraction kernels for each component of the field in terms of Bessel functions:

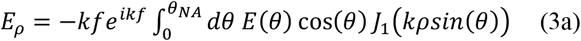

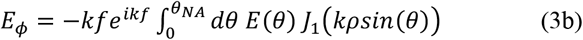

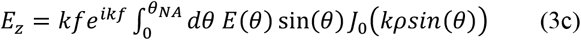

where *f* is the focal length of the lens, *ρ* is the radial coordinate at focus, and *θ*_*NA*_ is the maximum polar angle supported by the NA of the lens. From the solutions, it is evident that the RP and AP components have donut-like shapes governed by the first order Bessel kernel, whereas the ZP component will have a spot-like focus following the symmetry of the zeroth order Bessel function. While all polarization components at focus are shown in Eqs. 3a–3c for completeness, RP beams have explicitly zero azimuthal component at focus, and AP beams have neither RP nor ZP components.

In 2PM, the PSF is proportional to the square of the intensity of the focused light. In the case of an RP beam, the PSF includes orthogonal polarization components (Fig. 2b) which have a finite probability of interacting in an absorption event. This probability is dependent on the properties of the absorbing molecule (28), but here, we have assumed that the strength of this orthogonal absorption is equivalent to the co-polarized case; *i*.*e*. the PSF, *F(ρ)*, is given by *F*(*ρ*) = (|*E*_*ρ*_ |^2^ + |*E*_*z*_ |^2^)^2^. This assumption represents the most conservative estimation, as the interaction of the RP and ZP fields serves to widen the PSF and reduce resolution.

The relationship between the aspect ratio of the RP beam, *A*, and the PSF is shown in Figs. 2c–2e. As *A* increases, the strength of the RP component at focus decreases relative to the ZP component (Fig. 2c) corresponding to a decrease in the full-width-at-half-maximum (FWHM) of the PSF (Fig. 2d). However, this increase in lateral resolution is accompanied by a lengthening of the PSF in the axial direction (Fig. 2e), informing a tradeoff between lateral and axial resolution for different values of *A*. For the purposes of this work, we target an aspect ratio of *A* = 0.9 (red horizontal lines Figs. 2c–2e) as the resulting lateral resolution is comparable to a circularly-polarized Gaussian beam and there is minimal degradation in the axial resolution (green dashed lines Figs. 2d and 2e).

To facilitate SLAM, we must also consider the effect of the aspect ratio on AP beams (Fig. 3a). The electric field in this case is also described by Eq. 1 where the polarization vector has been changed from 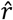 to 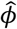.At focus, the PSF (Figs. 3b and 3c) is donut-shaped as expected, but has a second, weaker satellite ring surrounding the primary donut due to the increased aspect ratio of the beam. The width of the donut, *δ*, will ultimately inform the resolution enhancement, and, similarly to the FWHM of the PSF in the RP case, decreases with increasing *A* (Fig. 3d).

**Fig. 3:**
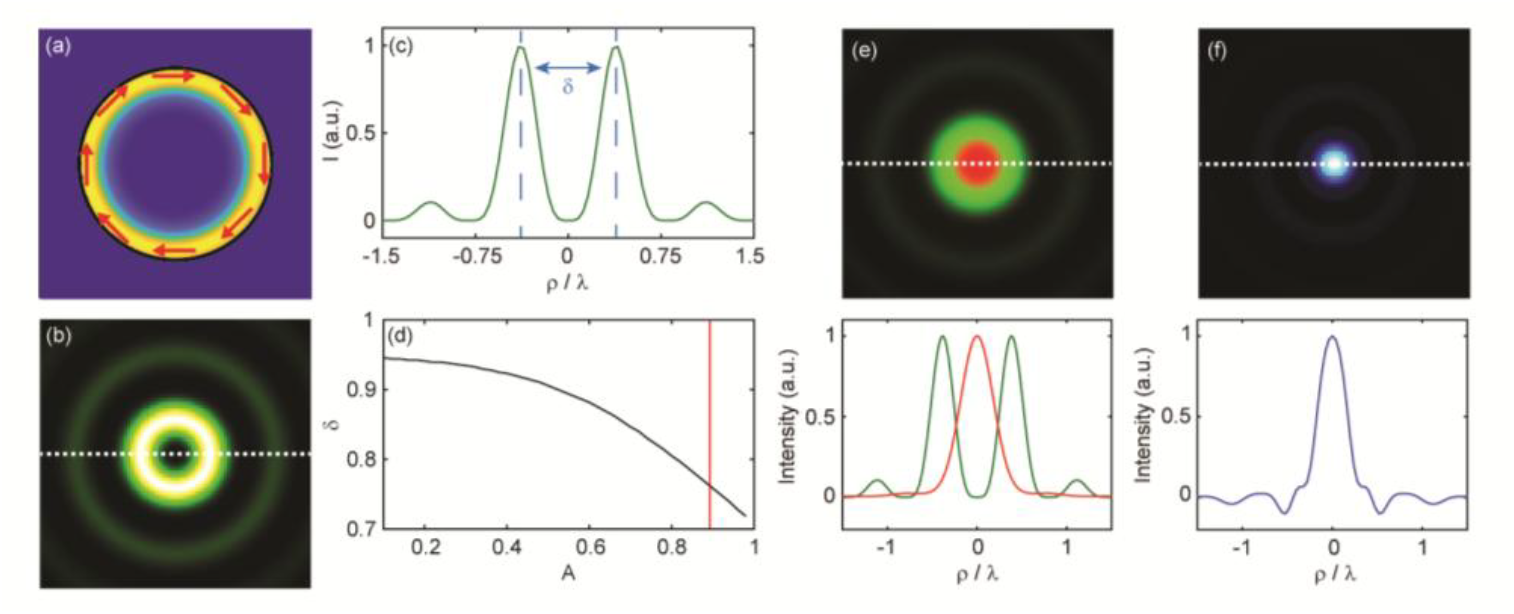
Annular AP beam in the far field (a), at focus (b), and in profile (c). (d) Beam width *δ* as a function of aspect ratio; aspect ratio for this work (*A* = 0.9) marked with red line. (e) Overlay of AP (green) and RP (red) beams at focus; line cut below. (f) Effective point spread function calculated by IWS; line cut below.

Figure 3e shows overlaid images of the RP and AP PSFs for *A* = 0.9; the overlap between the outer edge of the RP PSF and the inner edge of the donut suggests that resolution can be enhanced through subtraction. In SLAM, the super-resolved image is given by *I*_*SLAM*_ = *I*_*s*_ *– αI*_*d*_ where *I*_*s*_ and *I*_*d*_ are the images taken with the spot-like PSF and the donut-like PSF, respectively, and *α* is a weighting coefficient. Here, we employ intensity-weighted subtraction (IWS) to calculate our SLAM images (16). In IWS, *α* is spatially varying, rather than constant, such that *α(x,y) = [I*_*s*_*(x,y) – I*_*d*_*(x,y) +1]/2*, where *x* and *y* are the image coordinate vectors. In practice IWS has slightly less resolution enhancement relative to single-valued subtraction but reduces over-subtraction artifacts. The resulting IWS PSF (Fig. 3f) has a 1.2× smaller FWHM relative to the RP PSF and has minimal rippling or negative-valued artifacts, which are often apparent in conventional SLAM PSFs using single-valued subtraction.

## 3. Arbitrary beam shaping using MPLC

We employ MPLC to generate ring-shaped beams with arbitrary aspect ratios at the BFP of the objective. MPLC, as the name suggests, is a method for shaping an electric field distribution using multiple planes of phase modification (29–32). Here we apply MPLC to the most rudimentary form of the problem: high fidelity conversion of a single input field to a target output field using two phase plates. In this modality, the first phase plate controls the amplitude and the second controls the spatial phase, akin to classic phase retrieval systems (33,34). More generally, MPLC can employ many phase plates to map an ensemble of incoherent light beams into a set of co-propagating orthogonal spatial modes – of interest for mode division multiplexing schemes in telecommunications (30).

The crux of MPLC is the calculation of the necessary phase plates to perform the transformation. Here we employ the wavefront matching method to iteratively converge on solutions for each plate (29). In wavefront matching, an input Gaussian beam *E*_*i*_*(r)* impinges on the first phase plate, *M*_*1*_*(r)*, and propagates to the second plate resulting in the modified field 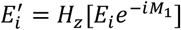 where *H*_*z*_ is an operator representing free space propagation over a plate-to-plate distance *z*. The difference in phase between the propagated field, *E*_*i*_′, and the target field, *E*_*t*_, informs an updated solution for the second phase plate:

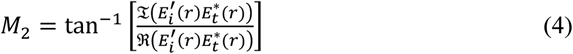

Similarly, the target field can be backward propagated, resulting in 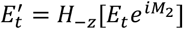 . The comparison of the modified target field with the input field, *E*_*i*_, then yields an updated solution for *M*_*1*_ following the same logic as Eq. 4. By iterating back and forth, the solutions for *M*_*1*_ and *M*_*2*_ converge, and the resultant phase plates can facilitate high fidelity conversion of the input beam to the target field.

## 4. Results

Figure 4a shows a schematic of our SLAM microscope. Light from an ultrafast laser (*λ* = 1035 nm, *τ* = 220 fs, *R* = 10 MHz, Thorlabs YFi-HP) is incident on a SLM (Hamamatsu X10468-07) with a two-bounce configuration to affect MPLC. The phase patterns *M*_*1*_ and *M*_*2*_ (inset Fig. 4a) are generated by our wavefront matching algorithm, which converges to a simulated 97.1% overlap with the desired *A* = 0.9 annular beam after 7 iterations. The output beam is generated at the surface of *M*_*2*_ and conjugated to the galvanometric scanners of our home-built two-photon microscope. A half-wave plate (HWP) is employed to set the polarization state to either vertical or horizontal and an annular aperture is used to remove artifacts in the center of the beam due to the finite conversion efficiency of the SLM. A vortex waveplate (VWP) is placed as close as possible to the BFP of the microscope objective (Nikon CFI75 Apo 25XC W, NA = 1.1). The size of the annular beam was chosen such that the total system magnification from *M*_*2*_ to the BFP (∼8×) ensures that the conjugated beam size matches the back aperture of the objective (*r*_*obj*_ ∼ 10 mm). An image of the annular beam at the BFP (inset Fig. 4a) shows azimuthal uniformity and confirms the desired aspect ratio. By imaging the same beam through a rotating polarizer (Fig. 4b), we confirmed that the polarization state is indeed RP with a polarization extinction ratio of >15 dB (97%) for horizontal polarization incident on the VWP.

**Fig. 4:**
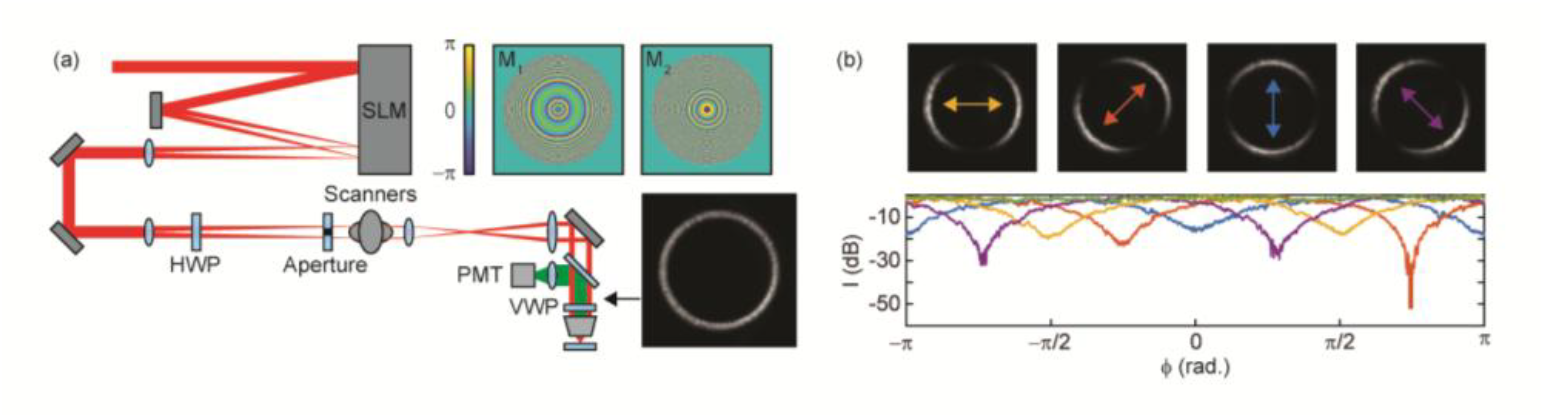
(a) Microscope schematic illustrating MPLC using a SLM programmed with phase patterns *M*_*1*_ and *M*_*2*_ (inset above), with plane *M*_*2*_ conjugated to the scanners of a conventional two photon microscope through an annular aperture to remove on axis zero order light; generated annular beam inset below; PMT = photo-multiplier tube. (b) Images of the annular beam through a polarizer at different angles (above) show azimuthal variation in intensity consistent with radial polarization (line plots below) with extinction >15 dB (97%) for all orientations.

To characterize the resolution of the microscope, we imaged 250 nm fluorescent beads (nile red). Figure 5 shows each of the constituent images: RP, AP, overlaid, and subtracted via IWS for an example measurement. As expected, the RP beam generates a spot-like PSF at focus with a FWHM of 460 ± 40 nm (mean and standard deviation of measurements of 32 beads determined by Gaussian fitting). For this microscope and objective, imaging with a Gaussian beam provides the same measured PSF width, 460 ± 40 nm, as the measurement for the RP beam – in line with the performance predicted by our simulations (Fig. 2d) and verifying that no resolution is lost by employing the RP beam for generating a spot-like focus. In the axial direction, the FWHM of the RP beam is 3.4 ± 0.1 μm, versus 1.7 ± 0.1 μm for the Gaussian beam, indicating a small reduction in resolution, also in agreement with our simulations.

**Fig. 5:**
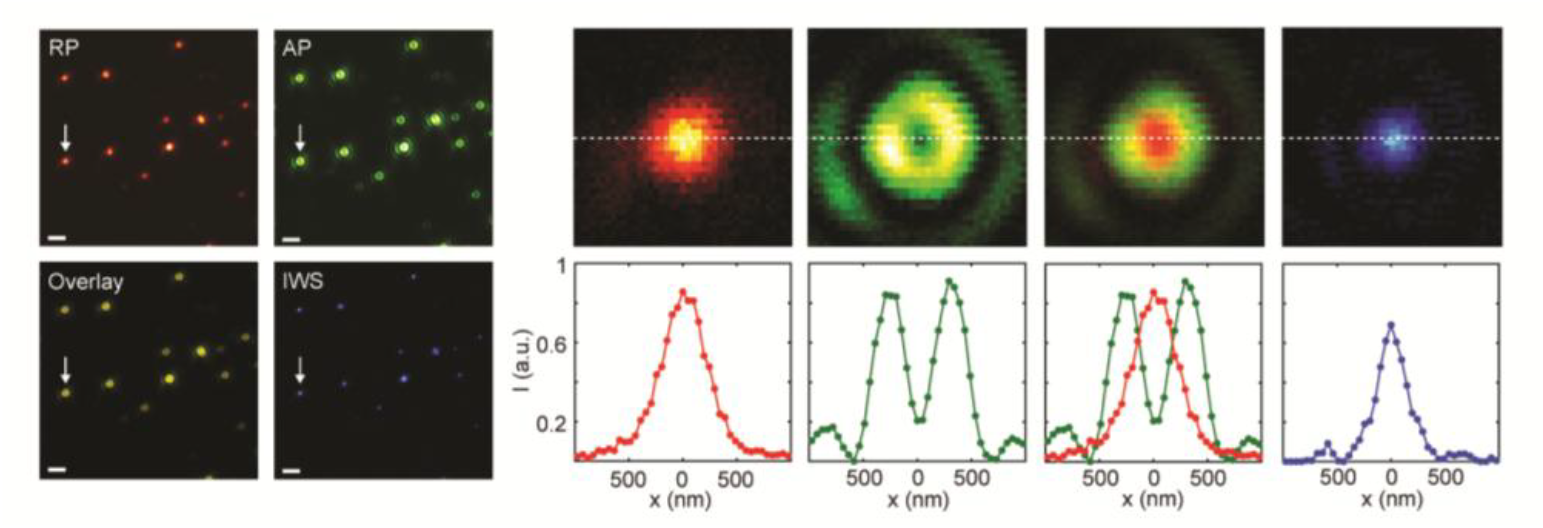
Fluorescent bead images using radially polarized (RP) and azimuthally polarized (AP) beams, and the resulting intensity weighted subtraction (IWS); zoom in and line cuts for beads marked by the white arrow are shown to the left; scale bar = 2 μm.

The AP beam forms the expected donut-shaped beam with surrounding ring. The width of the donut, *δ*, is 540 ± 40 nm – significantly narrower than the simulated value of 760 nm (Fig. 3a). We speculate that some of this increase in performance may be due to magnification mismatch generating a higher aspect ratio beam than intended at the BFP, however, surprisingly, we do not see similar improvement for the RP beam relative to simulations. This discrepancy may be due to the increased sensitivity of the RP beam to the exact incident beam geometry and polarization state, as the focal spot requires a favorable ratio of the ZP component relative to the residual RP light to form a tight focal spot. We also note that the AP width in our experiments is much narrower than would be obtained using a low aspect ratio, LG-like beam, as are commonly used in SLAM. Indeed, if we turn off our beam shaping, the donut beam width increases by a factor of ∼1.7.

As expected, the effective IWS beam has a tighter focus than the RP and Gaussian beams, with a FWHM of 300 ± 30 nm, 1.5 ± 0.1 -fold tighter than either single spot-like beam. This improvement out-performs the simulated expectation (1.2×), likely due to the narrower than expected width of the AP beam. However, 1.5× improvement in focal confinement is consistent with other reports in the literature for two-photon SLAM (11, 16). Additionally, artifacts outside of the central spot are minimal, owing to the use of the IWS algorithm, with only a single, weak ring visible in the image. Because the PSF of the AP beam has some residual intensity at its center, the signal at the center of the RP beam is necessarily reduced through subtraction; accordingly, the peak of the IWS beam is 0.8 ± 0.1 -fold smaller than the RP beam alone. Furthermore, we must consider that noise is additive in SLAM: assuming the same level of noise for the AP and RP beams, we estimate that the SNR is decreased by a factor of ∼1.8 through the subtraction process.

While FWHM has become a proxy for resolution as a simplification of the Rayleigh criterion, it is strictly speaking only valid for PSFs resembling the Airy disk. Thus, to validate the resolution of our system, we considered more general resolution criteria and thus conducted both Fourier ring correlation (FRC) and image decorrelation analysis (35–38). In FRC, two images with statistically independent noise are compared. The Fourier transform of each image is computed and FRC curves are determined by calculating the correlation between the distributions for pixels within the perimeter of a circle of a given radius in the spatial frequency domain. For small radii, correlation approaches unity, as low spatial frequency components are shared between the images. However, as the radius increases, noise begins to dominate, decreasing the correlation.

Figure 6a shows an example FRC curve generated from bead images captured using our microscope. The correlation strength falls off slower for the IWS image (blue curve) compared to the RP image (red curve) indicating improvement in resolution. Using the conventional value of 1/7 as a threshold value (black line), the resolution improves from 420 ± 30 nm to 290 ± 20 nm, a factor of 1.4 ± 0.1 (mean and standard deviation of 5 images), through SLAM.

**Fig. 6:**
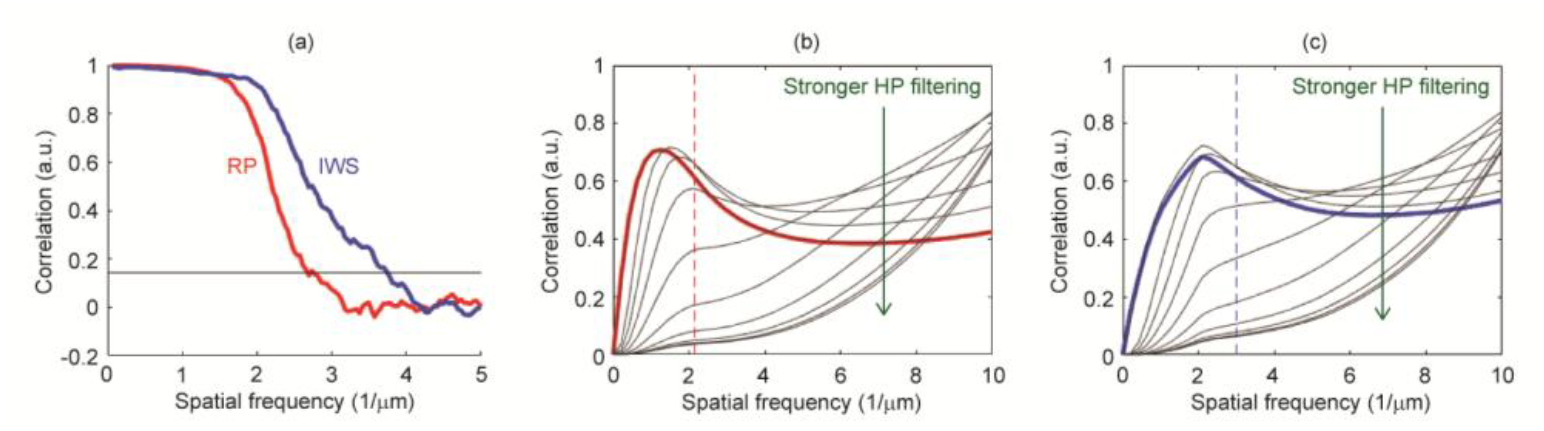
(a) Example Fourier ring correlation characterization of bead images for radially polarized (RP, red line) and intensity weighted subtraction (IWS, blue line) images; resolution threshold (1/7 = 0.143) shown by black line. Image decorrelation analysis for RP (b) and IWS (c) images; black lines denote correlation curves for various levels of high-pass (HP) filtering; dashed red and blue lines indicated predicted resolution.

In image decorrelation analysis, the Fourier transform of a single image is calculated, normalized, and filtered using a binary mask with a value of unity inside a given spatial frequency radius, and zero outside. Then the cross-correlation between the normalized, masked spatial frequency distribution with the unmodified distribution is calculated as a function of the radius of the mask, forming a correlation curve. For large radii, the correlation is less than unity due to the inclusion of both signal and high frequency noise contributions, but as the radii shrinks, the noise is preferentially filtered, resulting in an increase in correlation. At some point, the mask becomes small enough that signal is removed faster than noise, and the correlation decreases toward zero. Next, the same correlation curve calculation procedure is employed, but the initial image is subjected to high pass filtering with increasing strength. For weak filtering, the resulting curves resemble those of the non-high-pass filtered image, whereas stronger filtering eventually removes the peak in the correlation curve. Resolution is defined as the spatial frequency corresponding to the peak in the correlation curve for the strongest high-pass filtering where a peak is still evident.

Figures 6b and 6c show the results of decorrelation analysis on example bead images for RP and IWS beams, respectively. In each case, the non-high-pass filtered correlation curves are shown by the bold, colored lines, and the black lower weight lines are for high-pass filtered images. The decorrelation analysis predicts slightly lower resolutions for both the RP and IWS beams, 490 ± 30 nm and 370 ± 60 nm, than the FRC method, however, the improvement factor, 1.4 ± 0.3, is the same, indicating consistent improvement in resolution using our microscope.

## 5. Conclusions and outlook

We have demonstrated a SLAM microscope in a single-aperture, inline configuration. Our approach uses a single device to generate colinear RP and AP beams which in turn generate spot-like and donut-like PSFs under high NA focusing. Using MPLC, we optimize the far field electric field distribution at the BFP of the objective to ensure no loss in lateral resolution for the RP beam relative to imaging with a conventional Gaussian beam. Characterization of our microscope using fluorescent beads indicates an improvement in focal confinement by a factor of 1.5× relative to imaging with a Gaussian beam, and through subsequent correlation analysis we determine that the resolution of the microscope is as fine as 290 nm, or 0.28*λ*.

Crucially, this method is advantageous relative to conventional SLAM due to the consolidation of the system to a single beam path, removing the need for painstaking alignment of the donut and Gaussian beams, and eliminating the possibility of differential drift and subsequent registration artifacts. Our beam shaping methods ensure that the spot-like beam has the same resolution as a conventional Gaussian beam, despite passing through a VWP, and furthermore, our donut beam is 1.7× narrower than an unshaped beam – like those typically used in SLAM. Despite breaking the diffraction barrier, the resolution does not reach the nanoscale, as achieved by other super-resolution methods such as STED. However, in contrast to these methods, it requires lower laser power to attain super-resolution (39), works with conventional fluorescent labels, and has the potential to image both deep in tissue and *in vivo*. We expect that for applications requiring modest resolution enhancement, the relative simplicity, inherent 2PM compatibility, and deep imaging potential of our method would prove beneficial relative to more complex, depth-limited, super-resolution techniques.

In the current work, switching between the AP and RP beams is facilitated by mechanical rotation of a half-wave plate, slowing down the effective imaging rate. However, we intend to explore the use of electro-optic phase modulation as an alternative to mechanical rotation in future iterations of the microscope. Such modulators can operate with speeds faster than the repetition rate of our laser, allowing for switching between beams at the rate of pixel acquisition, and we expect that this increase in speed would help make the SLAM method robust to motion artifacts during *in vivo* imaging sessions. Furthermore, we expect use of a shorter wavelength laser would increase compatibility with conventionally used fluorophores as well as increase resolution without severe degradation to depth penetration: using an excitation wavelength of 900 nm, we expect the resolution would scale to ∼250 nm, and potentially even higher using post processing (18) or enhanced detection methods (40,41). Therefore, we expect this SLAM system could be an ideal candidate for super-resolution imaging experiments in thick, living tissue.

## Data availability

Data underlying the results presented this paper are not available publicly at this time but may be obtained from the authors upon reasonable request.

